# Females extract multiple signals from male song in Java sparrows

**DOI:** 10.1101/2024.02.08.579444

**Authors:** Rebecca N. Lewis, Hiroharu Makioka, R. Tucker Gilman, Masayo Soma

## Abstract

Songs play an important role in mate choice for many bird species. Females of some species prefer mates with song characteristics such as complexity, or with specific temporal or note features. In some species, females also prefer familiar songs, or songs that are acoustically similar to those of their fathers. These different aspects of songs may provide distinct information about the birds that produced those songs. For example, perceived familiarity may be a signal of local adaptation, locally relevant information or learned behaviours, or genetic compatibility, while song complexity may be a signal of overall genetic quality or ability to provide parental care. As a result, multiple axes of variation could influence female song preferences. We examined female preference for song familiarity and complexity using the Java sparrow (*Lonchura oryzivora*) as a model species. Females were presented with two pairs of song stimuli. In one pair one song was from the female’s father or from a social relative with an acoustically similar song and the other was unfamiliar. In the other pair the songs differed in two aspects of complexity: note repertoire and linearity. Females preferred their fathers’ songs over unfamiliar songs, but we did not find strong evidence that females generalized this preference to other socially related individuals. Females also preferred complex songs, specifically those with larger note repertoires and lower linearity. Our results suggest that female song preference in Java sparrows is multi-faceted, and females may obtain multiple kinds of information from male song.

## Introduction

Birdsong can provide a wide range of information about the individual producing it. As a result, songs often play an important role in female preferences and mate choice. Songs can provide information about overall male quality (Gil & Gahr, 2002) and about mate compatibility with the choosing female (Puur0nen et al., 2009; Searcy & Andersson, 1986). To convey this information, songs may vary along multiple axes, each providing different information to female receivers. Among female songbirds, preferences have been reported for a range of song features, including song familiarity (Fujii & Okanoya, 2022; Hernandez et al., 2009), complexity (Vyas et al., 2009), temporal characteristics (e.g., long, fast songs (Nolan & Hill, 2004)), and spectral characteristics (e.g., Tomaszycki & Adkins-Regan, 2005; Woolley & Doupe, 2008). How these features interact and contribute to overall song preference is likely to be complex. A preference for one song characteristic may overwhelm a preference for another, or multiple features may be evaluated and weighted independently (Dunning et al., 2014). Furthermore, preferences and weighting may vary among species and even among individuals.

Across species, two song features that often receive attention are song familiarity and song complexity. Song familiarity can provide females with information on mate compatibility at multiple scales. At the broadest scale, song familiarity can indicate individuals from the same (sub)species (Clayton, 1990). Within species, familiar songs may help to identify males from the same population that may be locally adapted (Podos & Warren, 2007) or have learned to exploit the local environment (Slagsvold & Wiebe, 2018; Wiebe, 2020). In line with this, females often show song preferences based on subspecies or macrogeographic population dialects, preferring songs from the subspecies or dialect they were exposed to early in life (Clayton, 1990; Hernandez et al., 2009). One mechanism by which these preferences could arise is sexual imprinting, where characteristics of the parental song phenotype are learned and used as a model for later preferences (ten Cate & Vos, 1999). Due to imprinting, females may have finer scale preferences within populations, for example, for their fathers’ songs or for songs that closely resemble those of their fathers. In zebra finches, preference for fathers’ songs is often reported, and there is some evidence that birds can generalize this preference to other similar songs within the population (Clayton, 1988; Miller, 1979; Wang et al., 2022; Wei et al., 2022). However, evidence for song preferences and generalization at this level remains limited in other species (e.g., Fujii & Okanoya, 2022)).

Complex songs, on the other hand, could signal male quality as opposed to compatibility. Complex songs may incur neural costs during development (Gil & Gahr, 2002). This link between song complexity and quality is supported by the negative influence of developmental stress (e.g., nutritional stress, parasite load) on song complexity in some species (Garamszegi, 2005; Soma et al., 2006; Zann & Cash, 2008). Complex songs may also indicate ‘good genes’ which can be passed on to offspring. In sedge warblers (*Acrocephalus schoenobaenus*), post-fledging survival of young is correlated with the genetic father’s but not the social father’s repertoire size (Hasselquist et al., 1996). Genetic background and environmental factors may also interact to influence song complexity (gene x environment interactions). For example, in zebra finches genetic background affected song complexity differently depending on nutritional regime (Woodgate et al., 2014). In addition, song complexity may also be related to the current condition of the signaller, for instance, increases in complexity with age are reported in some species (Kiefer et al., 2006; Kipper & Kiefer, 2010) or their capacity for parental care (Bartsch et al., 2015). Due to the potential individual and combined influences of early-life environment, genetics, and current condition, song complexity is likely to be an honest indicator of both good genes and the physiological capacity to provide parental care. Accordingly, preference for song complexity has been reported in some species (Hasselquist et al., 1996; Leitão et al., 2006).

We examined the roles of familiarity and complexity in song preferences using the Java sparrow (*Lonchura oryzivora*) as a model species. Each male Java sparrow learns to produce a single song during a critical period early in life. This song is used only in courtship (Ota & Soma, 2014), which also includes co-ordinated duet dancing in this species (Soma & Iwama, 2017). We used song playback trials to examine female preferences, which allowed us to test song preference independently of other male traits (Fujii et al., 2022). To determine females’ preference for familiar songs, we measured their responses to the songs of their fathers and of close relatives that sing similar songs, and we compared this with their responses to the songs of unrelated birds (hereafter, “familial” and “non-familial” songs, respectively). We expected that females would prefer their fathers’ songs, as predicted by sexual imprinting, and that females would be able to generalize this preference to the songs of other males that match their fathers’ song types, which would be important for avoiding close inbreeding during mate choice. To examine preference for song complexity, we presented females with songs that varied in note repertoire and song linearity (Scharff & No[ebohm, 1991). If complex songs are preferred, we expected that females would respond more strongly to songs with larger note repertoires and higher structural complexity (lower linearity). To our knowledge this is the first attempt to assess preferences for a song characteristic thought to indicate mate compatibility and another thought to indicate mate quality in the same study.

## Methods

### Subjects

Females were selected from a laboratory population of Java sparrows housed at Hokkaido University. Birds were housed at a temperature of 25 ± 3 °C, humidity 30 – 60%, with a 12L:12D lighting schedule. Birds were provided with seeds as food (a 3:1:1:1 mix of Japanese millet, fox tail millet, millet, and canary seed), shell grit, fresh green vegetables, and water ad libitum. Birds were usually housed in large single-sex cages (43 × 37 × 41 or 46.5 × 46.5 × 94 cm) with up to 20 individuals.

During breeding, birds were housed in family groups in cages (43 × 37 × 41 cm) that were visually but not audially isolated from other birds. Nests were inspected regularly, and eggs were cross-fostered when multiple nests with eggs were available. In Java sparrows, males sing a single song type, which remains consistent over their lifetime (Ota & Soma, 2014). However, some variation with age, particularly in early life (between song crystallization and 1 year old) is reported (Ota & Soma, 2014). Each bird learns its song from a tutor during a sensitive phase, which, in captive conditions, is usually the social father (Lewis et al., 2023; Lewis, Soma, et al., 2021). As a result, each social paternal line has a distinctive song type that can be identified visually in spectrograms. For the purpose of this study, songs produced by an individual’s social father or by another bird in the same social line were considered as “familial” (i.e., produced by or acoustically similar to songs produced by the social father).

In total, 17 cross-fostered females raised in the lab population took part in trials examining preference for song familiarity. None of the 17 females were social siblings from the same clutch, but 2 shared the same social father. Two pairs of females were genetic siblings but were not socially related. Females were aged ∼3.8 – 10.7 years (average ∼6.1 years) at time of testing. Twenty-four females took part in trials examining preference for song complexity: 11 cross-fostered females raised in the lab population, 3 hand-reared females (no song experience during development), and 10 females sourced from local breeders (and thus with unknown paternal song types and song experience during development). None of the lab-reared females in the complexity trials were social siblings, although there were 5 pairs of genetic siblings. It was not possible to determine the genetic and social relationships within two groups of birds (one of 2 females and one of 8 females) of similar age sourced from local breeders. Females were aged ∼0.65 – 6.9 years (average ∼3.5 years) at time of testing.

### Note classification

Classification of notes was necessary to measure complexity and to assess similarity between familial songs used in the trials. Notes were classified within social lineages based on visual inspection of the spectrograms (see Figures 1 and 2). All classifications were made by a single observer, but classifications were repeatable between observers with >95% accuracy (Lewis, Soma, et al., 2021). Because notes were classified within lines, we could not assess the similarity of songs from different social lines. Note characteristics are learned from social fathers, so the characteristics of note types can diverge among lines, and classifying notes across lines requires using broader note categories (Lewis, Soma, et al., 2021). These broader note categories are too coarse to capture accurately the song complexity and similarity within lines.

**Figure 1:**
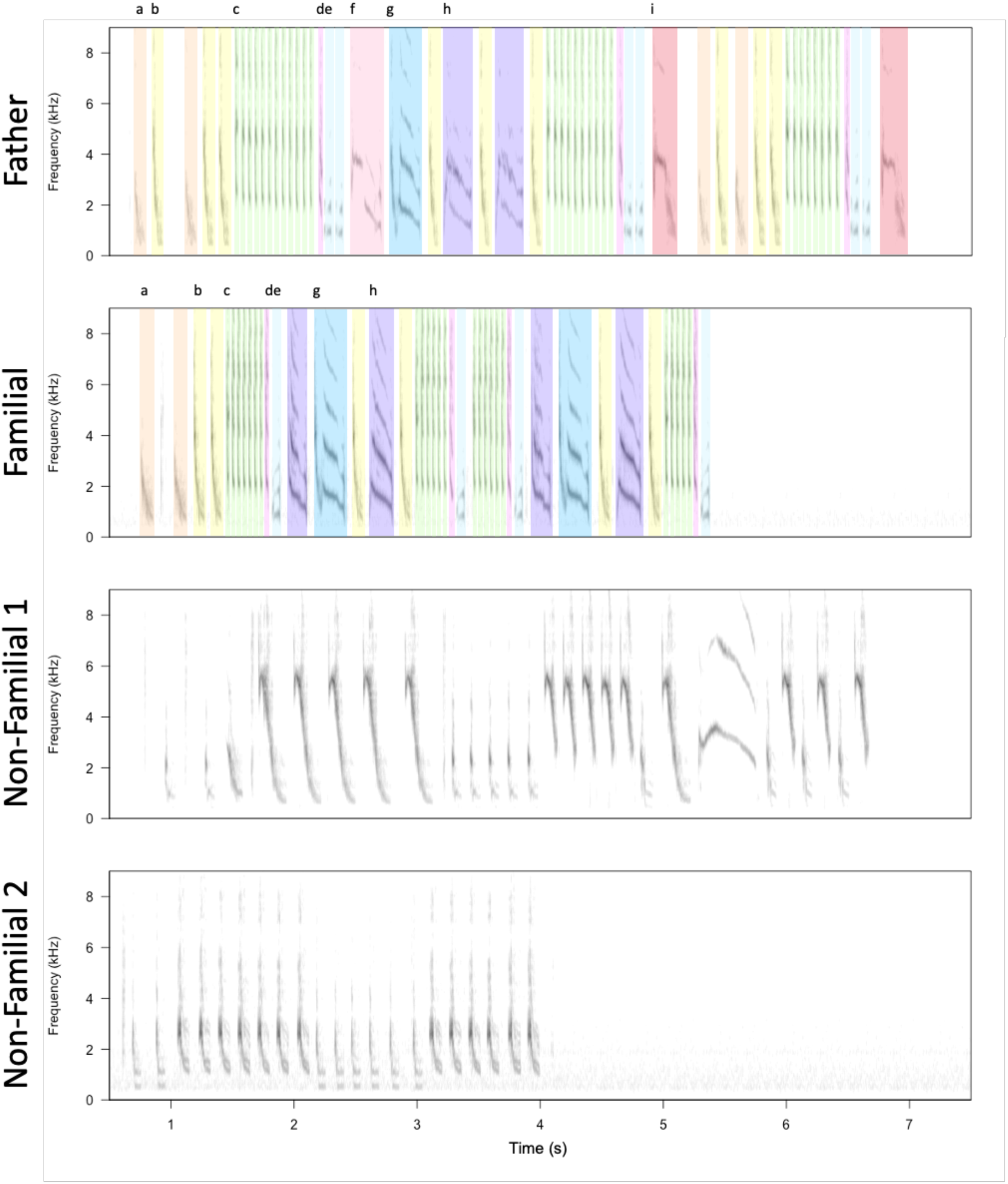
An example of stimuli presented to a single female in familiarity preference tests. The ‘Father’ and ‘Familial’ stimuli have been marked to show equivalent note types. Because notes from different lines are classified using different keys (i.e., different classification systems), notes in the unfamiliar songs are not assigned to types relevant to familiar songs. Spectrograms were created using Seewave (Sueur et al., 2008) (sample rate = 44.1kHz, window length = 512, overlap = 90%). X-axis scale (time (s)) is the same for all spectrograms.

**Figure 2:**
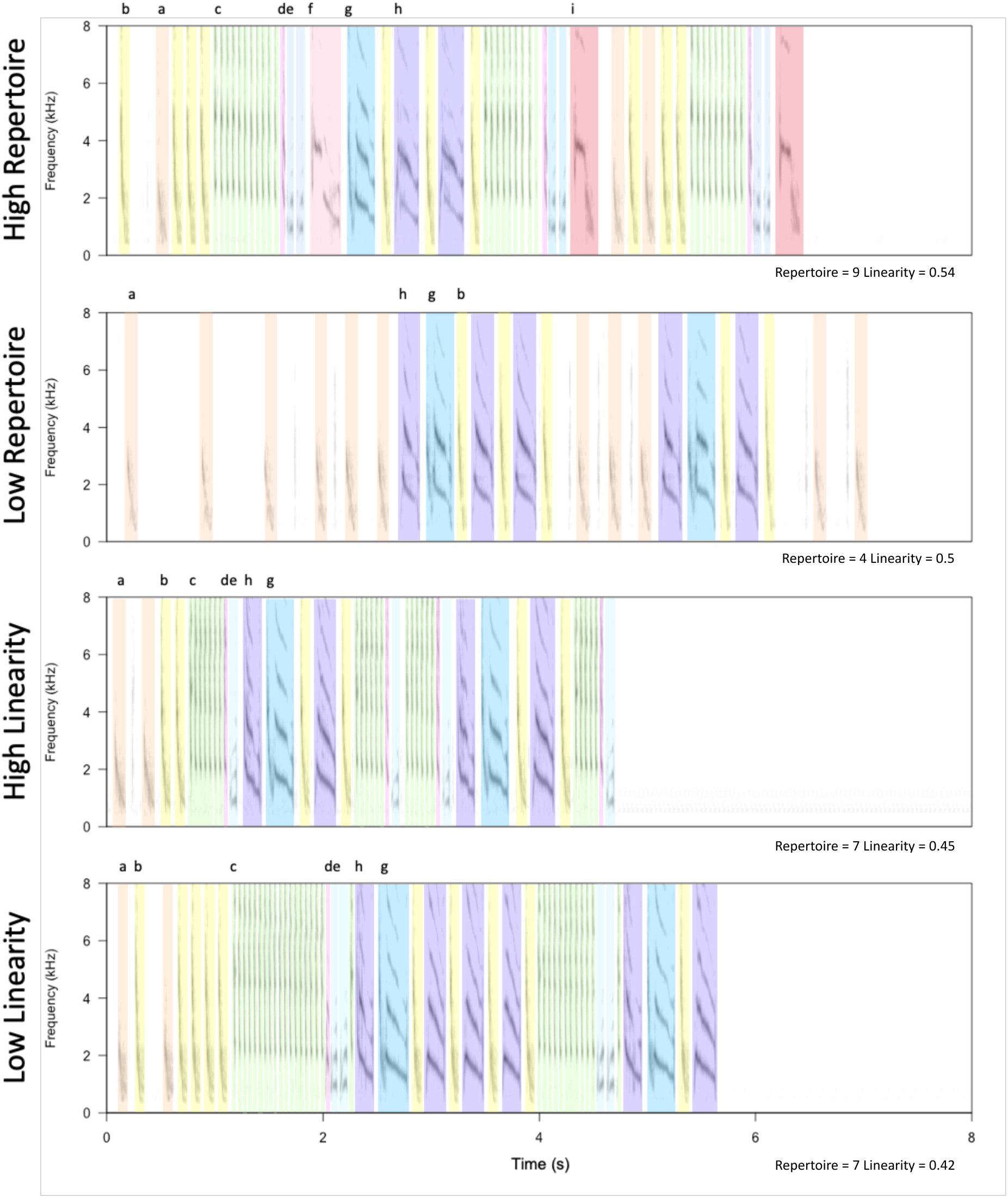
An example of stimuli presented to a single female during complexity preference tests. Stimuli have been marked to show equivalent note types. Spectrograms were created using Seewave (Sueur et al., 2008) (sample rate = 44.1kHz, window length = 512, overlap = 90%). X-axis scale (time (s)) is the same for all spectrograms.

### Stimulus assignments

#### Familiarity trials

Three types of stimuli were created for use in familiarity preference tests: ‘Father’, ‘Familial, and ‘Non-familial’ (Figure 1). Stimuli were presented to females in the study in pairs (see ‘Experimental trials’ below). For familiar songs, the ‘Father’ stimulus for each female was made up of songs recorded from the female’s social father (henceforth referred to as ‘father’). The ‘Familial’ stimulus for each female was made up of songs from a bird that had the same song type as the female’s father, i.e., from the same social line. Where fathers had been reared in the presence of subtutors (a non-father bird present during the sensitive period for song learning; n=4), visual inspection was used to confirm the father’s social line. In all but one case, the father’s (and therefore the female’s) social line was that of the father’s main tutor (i.e., the father’s social father). In one case, the female’s father’s song closely resembled his subtutor’s song, and so the father and female were assigned to the subtutor’s social line. Across familiar pairs, an average of 91.4% of father’s notes were also present in the ‘Familial’ stimulus (interquartile range 81.8-100%; range 57.1-100%) and an average of 72.7% of notes appeared in both the ‘Father’ and ‘Familial’ stimuli (interquartile range 58.6-93.8%; range 26.8- 100).

Each female was assigned two ‘Non-familial’ stimuli to prevent habituation to a single non-familial song. Familial and non-familial birds were selected at random for each female. Where possible, clutchmates were avoided, but in two cases no other familial birds were available. Analyses were conducted with and without these trials included, and the removal of these two cases did not qualitatively affect our results (Supplementary Material 1). Stimuli were constructed using songs from 35 males. Many males appeared as both familial and non-familial birds in the stimuli (11 father/familial only, 7 non-familial only, 17 father/familial to some females and non-familial to others) Each stimulus was heard by a median of 2 females (range 1 – 5).

#### Complexity

We used two measures of complexity to assess females’ preference for complex songs: note repertoire and song linearity. Note repertoire is a measure of note diversity, defined as the number of unique note types used by a bird within their song, with a high note repertoire associated with complexity. In this study, a song’s note repertoire was taken as the number of unique note types within a song and was averaged across the three songs used to construct the stimulus to give a single value. Linearity is a measure of structural complexity of a song, defined as the number of unique note types divided by the number of unique note-to-note transitions, with low linearity associated with high complexity (Scharff & No[ebohm, 1991). As with repertoire, linearity was calculated per song and averaged across songs used within a stimulus to give a single value. Four types of stimuli were created for use in complexity preference tests, ‘High note repertoire’, ‘Low note repertoire’, ‘High linearity’, and ‘Low linearity’, which were presented in ‘High’ vs. ‘Low’ pairs for each measure. Males for each stimulus set in the complexity trials were selected from the same social line to control for potential preferences among females for songs from specific lines. The differences between ‘High’ and ‘Low’ in each category varied among social lines based on individual variability and the availability of archival recordings. For each stimulus, females heard three different songs from the same male (see Stimulus Construction). Across all stimulus pairs, the mean difference in repertoire was 2.33 notes (median = 1.67, range = 0 – 5) and the mean difference in linearity was 0.061 (median = 0.054, range = 0 – 0.207). Stimuli contained natural songs, which were unmanipulated in terms of their features. As a result, whilst pairings were chosen based on complexity, other song features also varied across the songs.

Each female was assigned a set of four stimuli from the same social line (example set in Figure 2). This controls for possible differences in attractiveness between the song types of the different social lines. Where females were reared by parents in the main population, their assigned stimuli were drawn from their father’s social line (n=11). If females were hand-reared or were reared outside of the main population (i.e., birds sourced from local breeders), they were assigned a set of stimuli from available recordings belonging one social line (n=13). Where females received songs from their father’s social line, songs from the father or clutchmates were avoided where possible. However, as this could not always be avoided, the level of relationship was accounted for during data analysis. For one female, only three stimuli were available in the father’s social line, so one stimulus was repeated in each of the trial types, i.e., the low linearity and low repertoire stimuli consisted of the same songs.

### Stimulus construction

Stimuli were constructed using available archival recordings from individuals in the lab population. Archival recordings had 44.1 kHz sampling rate and 16-bit resolution and consisted of individual birds singing alone. Recordings were made using a digital sound recorder (Marantz PMD 661, Zoom Q3HD, TASCAM DR-100 MKIII). Recordings were high-pass filtered at 375 Hz in Raven Pro (1.6.3) to remove background noise. Recordings from all four of the birds from one of the social lines had been pre- processed with a high pass filter of 1 kHz. A high pass filter of 1 kHz is unlikely to have removed any important song features, and the closely related zebra finch (as well as other songbirds) are reported to have a narrow hearing range with highest sensitivity between 1-6 kHz (Hashino & Okanoya, 1989). Thus, we do not expect this to have an effect on preference tests.

For each stimulus, three songs from a single male were selected. The three songs were repeated at a constant rate (one song commencing every 10s) for two minutes, alternating among songs. Thus, a full stimulus contained 12 songs in the pattern ABCABCABCABC and lasted a total of 2 minutes. Many of the archival songs were recorded with recorders using stop start direction (so only significant noise events were recorded), or, had been segmented into individual songs prior to storage. As such, we did not have information on the natural song rates of individuals in our dataset, and there is little published information on song rates in this species. Since song rate can influence preference (Forstmeier, 2004; Nolan & Hill, 2004) we chose to control for song rate during playback. The chosen rate may differ from natural song rates, and natural song rates are likely to differ among individuals and social contexts, as in other species (e.g., zebra finch (Dunn & Zann, 1996; Jesse & Riebel, 2012; Naguib et al., 2008)). As song rate was held constant during stimulus construction, sound density (i.e., the total time occupied by songs within the stimulus) differed among stimuli where individuals had differing song lengths. We accounted for in this in analyses (see Data Analysis). Across all stimuli, amplitude was equalized by matching peak amplitude between stimuli, as song amplitude may influence preference (Ritschard et al., 2010)().

### Experimental trials

Except where explained below, experimental and equipment setup was the same for both familiarity and complexity trials. Females were transferred from their usual cage to a smaller, individual cage (37 x 25 x 19 cm) for the duration of the playback experiment. To complete individual trials, females were placed in a soundproof chamber. Whilst females were not completing trials or habituating to the soundproof chamber, they remained in their individual cages, but were not socially isolated from other birds. Food and water were provided *ad libitum* throughout the experimental period. Following completion of trials, females were returned to their usual cage.

For all trials, playback was conducted using a speaker (Micro wireless, JBL, California, USA) placed approximately 15 cm from the front of the cage. Amplitude was equalized across stimuli, and speaker volume was held constant across the experimental period, but the amplitude of playback within the soundproof chamber was not measured. Trials were recorded using a video camera (Q4n, ZOOM, Tokyo, Japan).

#### Familiarity

There were two types of trials during the familiarity preference tests, Father vs. Non-familial and Familial vs. Non-familial, with each pairing using a different non-familial stimulus. For two birds, recordings of the father’s song were not available, so two Familial vs. Non-familial stimulus pairs were presented, with a different birds in each pair.

Females completed four trials during the experiment, with each trial taking place on a different day (mean days between trials = 2.94, standard deviation = 1.19 range = 2 – 6). The order of the stimuli within pairs was reversed in successive trials (Figure 3), and the order of trials was varied among birds. Prior to each trial, females in individual cages were placed in a soundproof chamber and allowed to habituate to the environment overnight. At the start of each trial, following the chamber door being opened to switch on equipment to begin playback, females completed a short habituation period (5 minutes). After this, females were presented with four stimuli (2 minutes each), alternating between familiar and unfamiliar, with a 30 second gap between each stimulus presentation (Figure 3). Each trial lasted 10 minutes. One of the 17 birds was removed from the study after completing a single trial. In total, 65 trials (4 trials for each of 16 birds plus one trial for one bird) were recorded for analysis.

**Figure 3:**
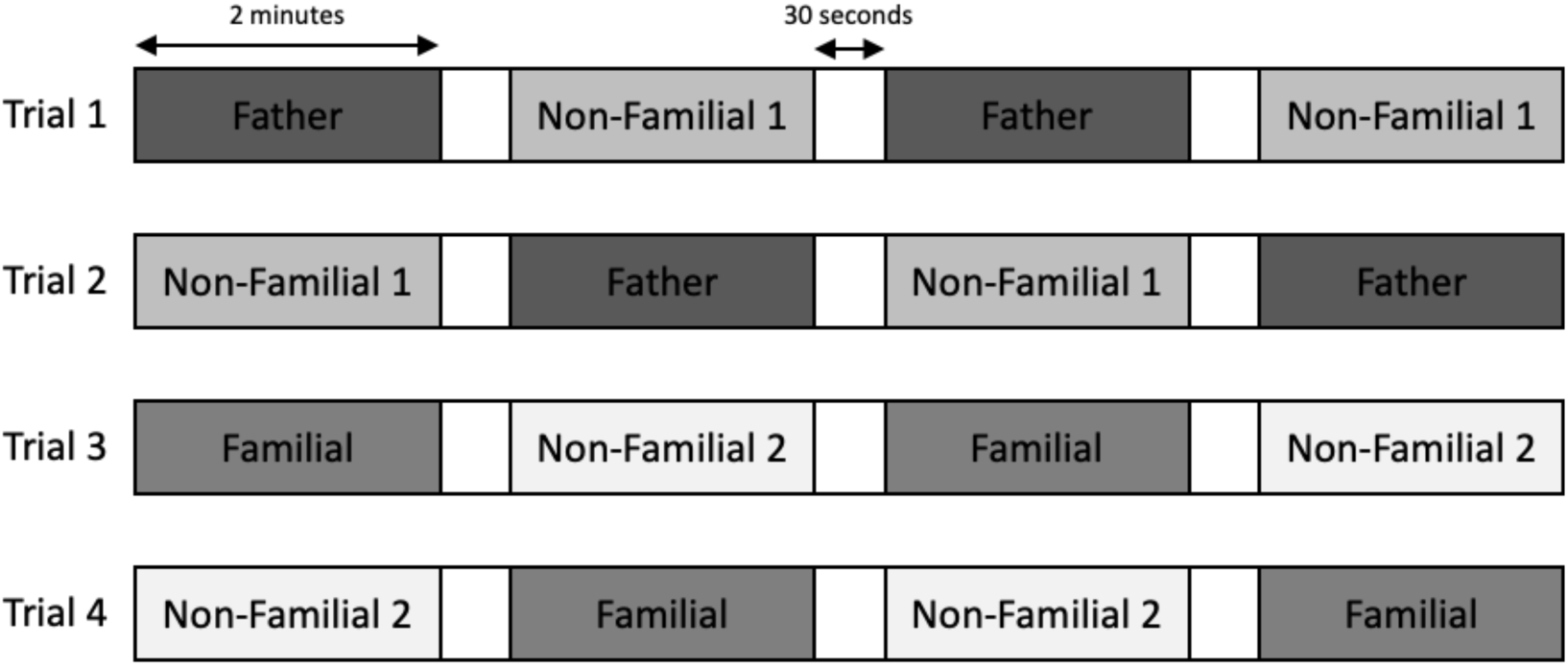
Example experimental set up for a female completing the familiarity preference tests. Order of trial presentation varied among females.

#### Complexity

Complexity preference tests also comprised two trial types, ‘high’ vs. ‘low’ note repertoire and ‘high’ vs. ‘low’ linearity. As with the familiarity preference tests, females completed four trials during the experiment and each trial took place on a different day (3-day gap between each trial). Each trial type was completed twice with the order of presentation reversed in each repeat (Figure 4), and the order of trials was varied among females. Trial presentation was identical to that for the familiarity preference tests, with a short habituation period (5 minutes) followed by the presentation of stimuli. Based on females’ responses in the familiarity preference tests, overnight habituation was not deemed necessary, as birds habituated quickly to the soundproof chamber. In total, 96 trials were recorded for analysis. One trial could not be analysed due to an equipment malfunction, so data was extracted from 95 trials.

**Figure 4:**
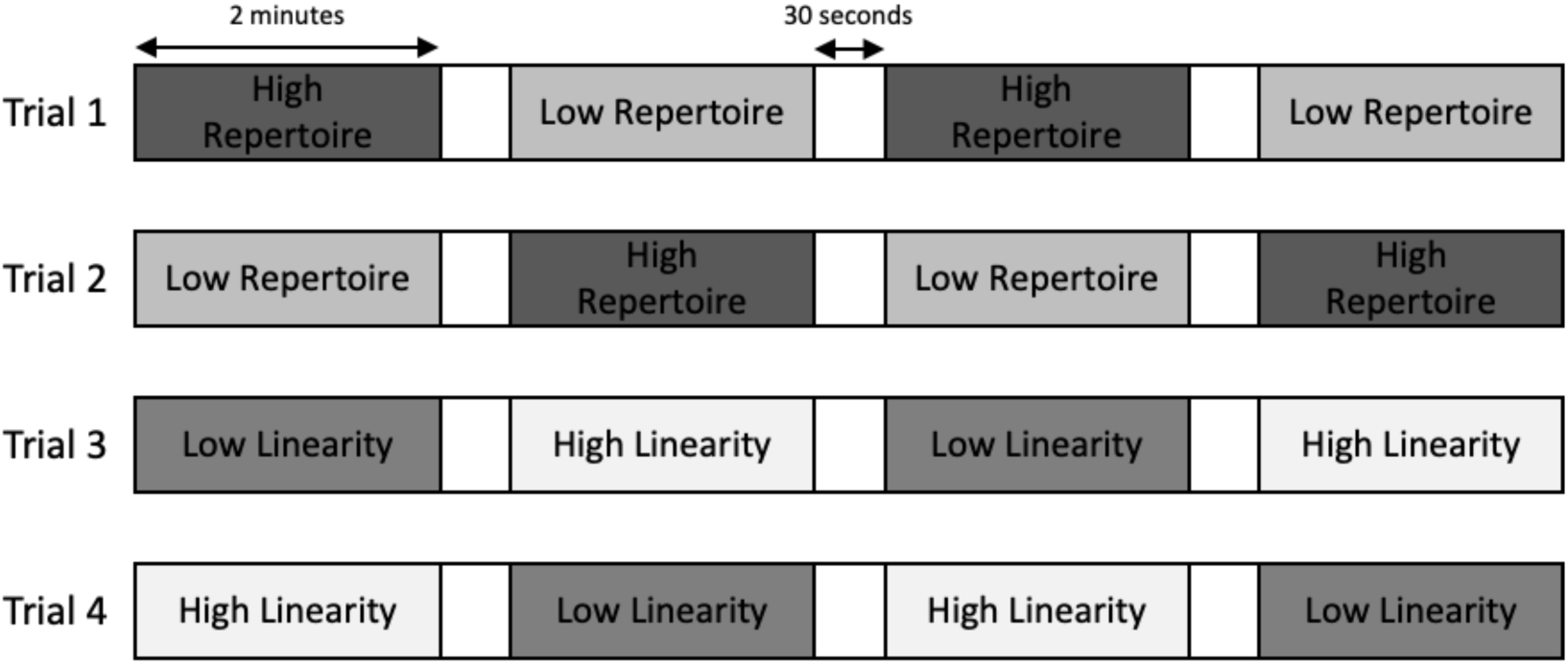
Example experimental set up for a female completing the complexity preference tests. Order of trial presentation varied among females.

### Behavioural data

The extraction of behavioural data was the same for both familiarity preference and complexity preference tests. Information on behavioural responses to stimuli was extracted from video footage taken during experimental trials. We examined four behaviours that are performed in a range of social situations by Java sparrows: calls, hops, bill wiping and fluffing. These behaviours were chosen to cover a wide range of social responses, including behaviours used in courtship displays, as these may indicate mate preference. Java sparrows produce contact calls, which consist of a single, discrete note, in social interactions (Restall, 1996). Whilst not explored in Java sparrows, in the closely related Bengalese finch, calls are a reliable indicator of mate preference, showing similar patterns to copulation solicitation displays (Dunning et al., 2014). In Java sparrows, both hopping and bill wiping are associated with courtship dancing (Soma & Iwama, 2017), so may indicate mate preference. In another Estrildid finch, the zebra finch, fluffing behaviour is associated with mate preference and is performed in response to complex, attractive songs (Vyas et al., 2009). Juveniles experiencing more fluffing behaviour in response to songs also developed more accurate songs (Carouso-Peck et al., 2020; Carouso-Peck & Goldstein, 2019).

Instances of each behaviour were counted for each stimulus presentation (i.e., each 2-minute block of song playback), and these values were used in the analysis presented in the main text. Motivations behind behaviours performed during the silent gaps may be hard to interpret. However, responses in the silent gaps were correlated with the number of responses in the preceding stimulus presentation for most behaviours examined. The inclusion of behaviours performed in the silent intervals did not qualitatively affect our results (Supplementary Material 2). Behaviours during the 5-minute habituation period were not counted, as birds tended to react as equipment was switched on and the door to the soundproof chamber was closed.

### Data analysis

All analyses were conducted in R (ver. 4.1.3) (R Core Team, 2022). Within each set of tests (familiarity preference tests and complexity preference tests), we conducted separate tests for each of calls, hops, bill wiping and fluffing. Due to the number of non-responses from females, it was possible that our data were zero-inflated. We initially fit Poisson regressions to the data and tested for zero-inflation using the performance package in R (Ludecke et al., 2021). Poisson error distribution was chosen as the data consisted of counts of behaviours within each trial. Where likely zero-inflation was detected, we re-ran the models including a zero-inflation term. As a result, some behavioural responses were analysed with generalized linear mixed effect models with Poisson error distribution using the lme4 and lmerTest packages (Bates et al., 2015; Kuznetsovs et al., 2017), and others were analysed using zero-inflated Poisson models using the glmmTMB package (Brooks et al., 2017).

Where models did not converge, we sequentially removed the random effect (from either the conditional or zero-inflation) that explained the least variance until the model converged. In all but one case (see Supplementary Information 2), the removed random effects explained less than 0.001% of the total variance in the model. Models for familiarity and complexity preference tests were broadly similar, but differed in some predictors, as outlined below.

#### Familiarity

For each model, the behaviour of interest was used as the response variable. We included the type of stimulus (familial or non-familial), the type of trial (Father vs. Non-familial or Familial vs. Non-familial), the order of the stimulus within the trial (played 1^st^, 2^nd^, 3^rd^ or 4^th^), and the order of the trial within the experimental period (1^st^ trial, 2^nd^ trial, 3^rd^ trial or 4^th^ trial) as fixed effects in the model. The order of the stimulus was included to account for habituation within the 10-minute trial. The order of the trial was included to account for habituation over the course of the overall experiment. We included the average song duration (start of first note to end of last note (s)) for each stimulus. This accounts for the differences in song length between individuals, as song duration (or increased sound density) may affect females’ responses (Clayton & Pröve, 1989; Neubauer, 1999). We also included the interaction between the type of stimulus and the type of trial to determine if the difference in response to familiar songs was different in Father vs. Non-familial or Familial vs. Non-familial trials.

We included the female ID, the ID of the male that produced the stimulus song and female ID x male ID as random effects in the models. Where used (calling and hopping behaviour), the zero-inflation formula included an intercept and a random effect of the individual female. This accounted for the fact that females may not respond during the presentation of a stimulus during the trial, and that the likelihood of not responding differed among females. Zero inflation models with additional fixed (familiar vs. unfamiliar, test type) and random (male ID, female ID x male ID) effects did not improve model fit or alter qualitative results.

#### Complexity

As with familiarity preference tests, the behaviour of interest was used as the response variable. We included the average note repertoire and average linearity of the stimulus as fixed effects. Complexity measures were included as numerical values, rather than ‘high’ or ‘low’. This is because the difference between the ‘high’ and ‘low’ values differed depending on the social line from which the stimuli were selected. As with the familiarity preference tests, we included the order of the stimulus within the trial and the order of the trial within the experimental period as fixed effects to account for habituation at different scales. As some females heard stimuli from their father’s social line (n=11), which may have included their father or cluthmates, we also included fixed effects of stimulus familiarity (familial vs. non-familial) and of relatedness (father, clutchmate, or unrelated) to account for this, as birds may respond differently to familiar stimuli or stimuli from close relatives compared to unfamiliar or unrelated stimuli. As with familiarity, we included the average song duration for each stimulus. As hand-reared birds had less experience with adult song models, their responses and preferences may be different to parent reared birds. To account for this, hand-rearing status was included as a fixed effect in the models. We included the same random effects as in the familiarity models: female ID, the ID of the male that produced the stimulus song and female ID x male ID. Where used (calling, hopping, and bill wiping behaviour), the zero-inflation formula included an intercept, fixed effects of repertoire and linearity, and random effects of female ID, the ID of the male producing the stimulus song, and female ID x male ID.

### Data availability

Data and code associated with this study can be found on Figshare: https://doi.org/10.48420/25142780.v3.

### Ethical approval

This study was approved by The University of Manchester Animal Welfare and Ethics in Research Board (Permit Number D0062) and Hokkaido University (Permit Numbers 16-0020 and 22-0052).

## Results

### Familiarity

Collinearity between behavioural responses varied between 0.087 and 0.431 (Supplementary Table 3A). Overall, we found evidence that Java sparrow females responded more strongly to their fathers’ songs than to non-familial songs (Table 1, Figure 5). Significantly more calls, hops and marginally more fluffs were performed during playback of fathers’ songs. The number of bill wipes followed the same trend but did not reach marginal significance. Females did not show significantly stronger responses to ‘Familial’ songs than to non-familial songs.

**Figure 5:**
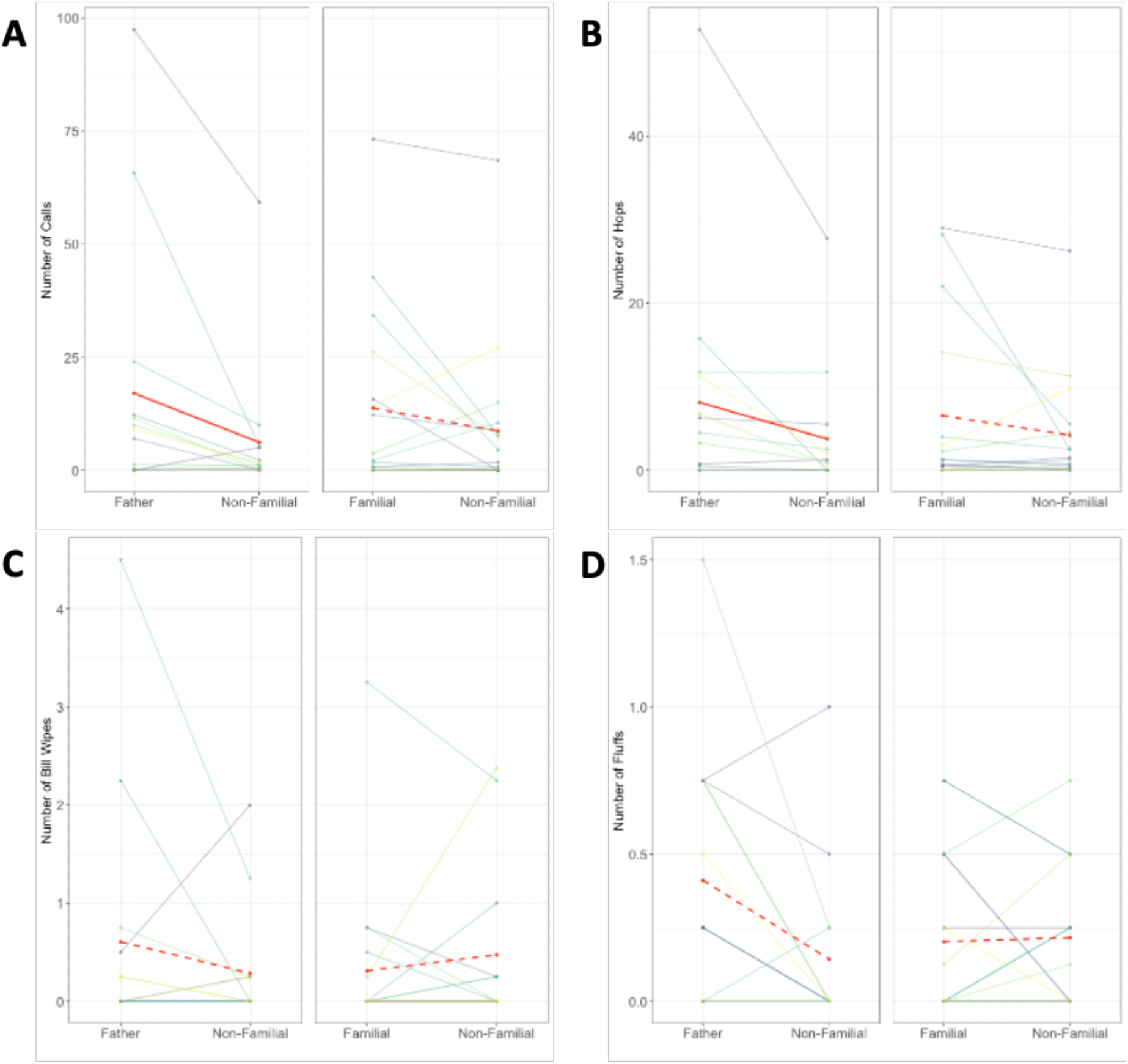
Behavioural responses from females to familial and non-familial songs in Father vs. Non-familial and Familial vs. Non-familial trials: A) calls B) hops C) bill wiping D) fluffing. Number of responses is averaged across stimuli/trials of the same type. Transparent lines represent individual females’ responses. Bold red lines indicate the average response across individuals within the trial type. Significant differences are indicated by solid lines, nonsignificant differences are indicated by dashed lines.

**Table 1:** Results of mixed effect models for behavioural responses to song familiarity. Estimates show the effect of the predictor on the logged number of responses. Statistical models included an interaction term involving test type (Father vs. Non-familial or Familial vs. Non-familial) and song familiarity (Familial vs. Non-familial). For ease of interpretation, effect sizes and p-values have been calculated from the model output for each comparison.

|  | Estimate | p-value | Direction of effect |
| --- | --- | --- | --- |
| **Number of Calls** |  |  |  |
| **Father vs. Non-familial** | **0.753** | **0.015** | **Greater response to father's songs** |
| Familial vs. Non-familial | 0.328 | 0.201 | N.S. |
| **Stimulus order** | **-0.215** | **<0.001** | **Habituation effect** |
| **Test Order** | **0.176** | **<0.001** | **Acceleration effect** |
| **Average Duration** | **0.291** | **0.011** | **Greater response longer songs** |
| **Number of Hops** |  |  |  |
| **Father vs. Non-familial** | **1.335** | **<0.001** | **Greater response to father's songs** |
| Familial vs. Non-familial | 0.475 | 0.144 | N.S. |
| **Stimulus order** | **-0.089** | **<0.001** | **Habituation effect** |
| **Test Order** | **-0.107** | **<0.001** | **Habituation effect** |
| Average Duration | -0.038 | 0.723 | N.S. |
| **Number of Bill Wipes** |  |  |  |
| Father vs. Non-familial | 0.332 | 0.641 | N.S. |
| Familial vs. Non-familial | -0.518 | 0.355 | N.S. |
| Stimulus order | 0.089 | 0.310 | N.S. |
| **Test Order** | **0.348** | **<0.001** | **Acceleration effect** |
| Average Duration | 0.123 | 0.503 | N.S. |
| **Number of Fluffs** |  |  |  |
| *Father vs. Non-familial* | *0.764* | *0.088* | *N.S. (Direction of greater response to father's songs)* |
| Familial vs. Non-familial | 0.015 | 0.969 | N.S. |
| **Stimulus order** | **-0.275** | **0.019** | **Habituation effect** |
| Test Order | 0.012 | 0.922 | N.S. |
| **Average Duration** | **0.250** | **0.024** | **Greater response to longer songs** |

Females performed more calls and fluffs in response to stimuli with higher sound density (longer songs) (Table 1).

Birds responded with less calling, hopping, and fluffing behaviour to stimuli presented later in trials (Table 1), suggesting habituation over the course of 10-minute trials. Changes in behaviour over the course of the whole experiment were less clear-cut (Table 1). Whilst there was a decrease in hopping behaviour across experimental trials, this was not the case for other behaviours. We found no significant differences in fluffing behaviour across the course of the experiment. For calling and bill-wiping behaviour, we found evidence for an acceleration effect; birds showed greater responses in later trials. This may suggest a change in preferred response type over the course of the experiment.

### Complexity

Collinearity between behavioural responses varied between 0.092 and 0.381 (Supplementary Table 3B). Overall, Java sparrow females preferred songs with larger note repertoires (Table 2; Figure 6). Females were more likely to call (B=0.453, p=0.032) and produced more fluffing behaviour (B=0.140, p=0.043) when stimuli included more note types. The numbers of hops and bill wipes were directionally similar but did not reach marginal significance. The effect of linearity was less pronounced (Table 2, Figure 7). Females were less likely to call in response to more linear (i.e., simpler) songs (B=-15.477, p=0.026), but linearity did not predict the frequency of any other response type. Females called more in response to stimuli with greater sound density (longer songs), but significant patterns were not apparent for other behavioural responses.

**Figure 6:**
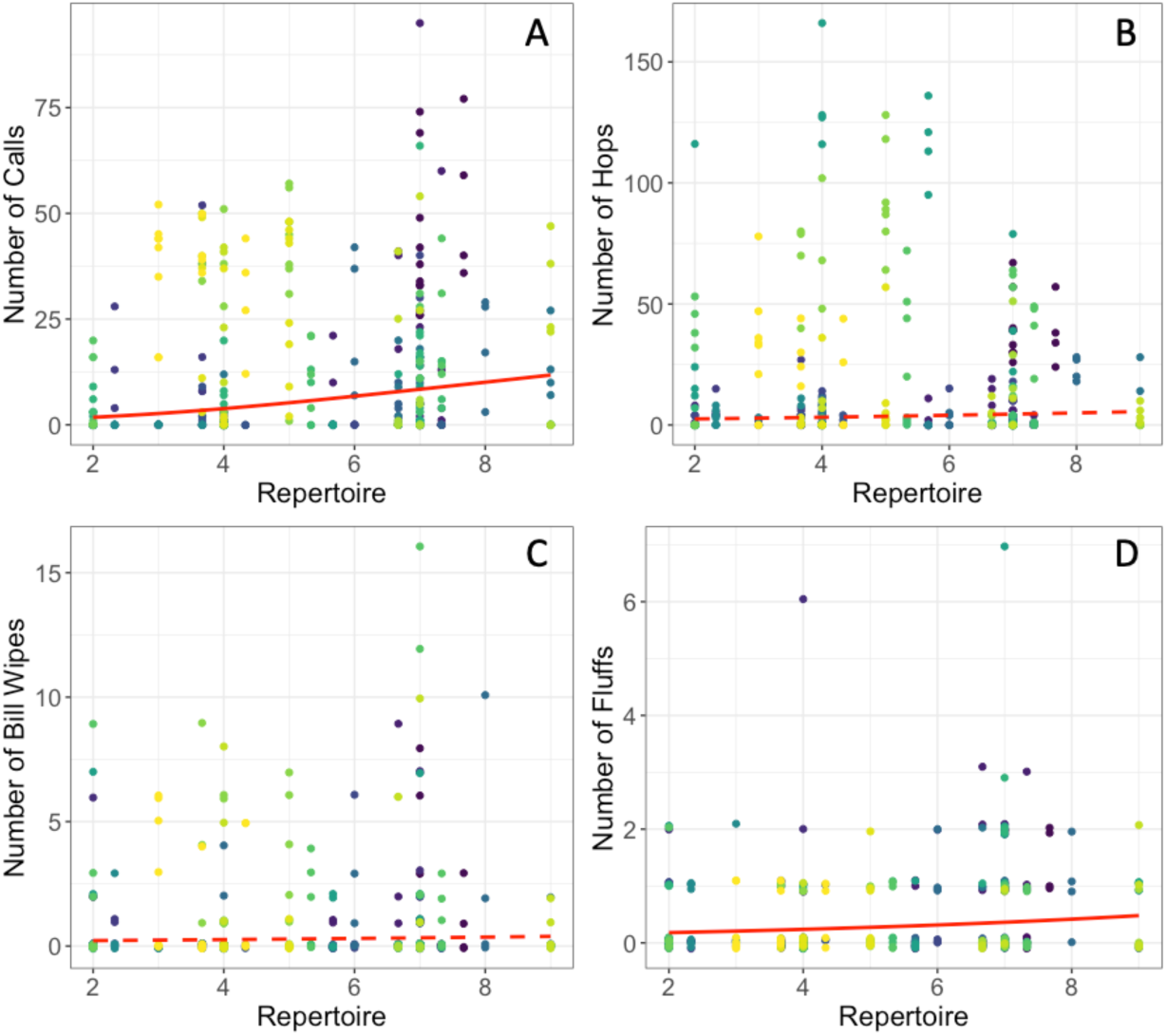
Behavioural responses from females to stimuli in complexity preference tests compared to stimulus repertoire: A) calls B) hops C) bill wiping D) fluffing. Coloured points represent individual females’ responses to stimuli. Bold red lines show the model predictions for each behaviour, significant results are represented by solid lines, nonsignificant results are represented by dashed lines.

**Figure 7:**
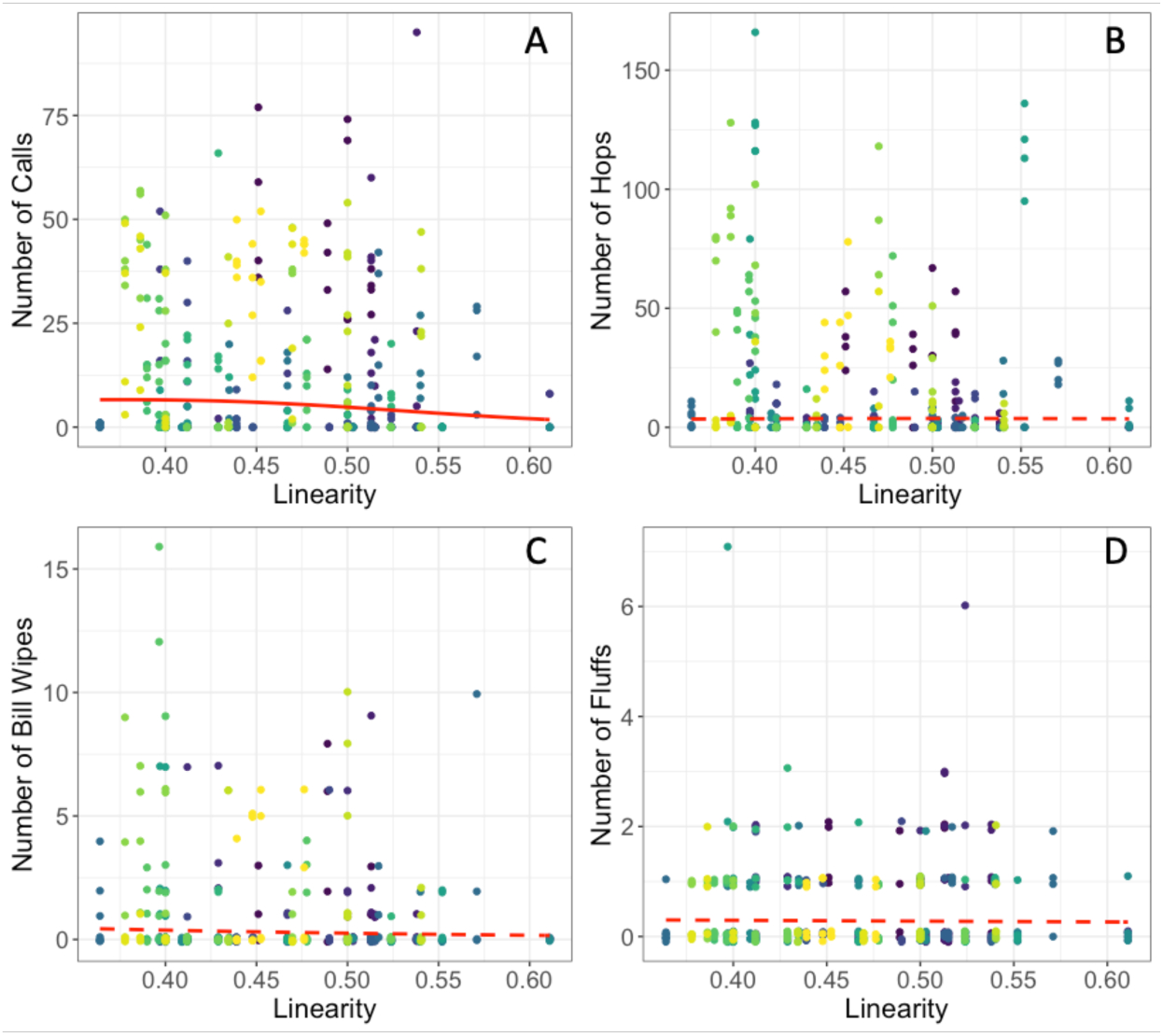
Behavioural responses from females to stimuli in complexity preference tests compared to stimulus linearity: A) calls B) hops C) bill wiping D) fluffing. Coloured points represent individual female’s responses to stimuli. Bold red lines show the model predictions for each behaviour, significant results are represented by solid lines, nonsignificant results are represented by dashed lines.

**Table 2:** Results of mixed effect models for behavioural responses to song complexity. Estimates in the top (or only) row of each cell show the effect of the predictor on the logged number of responses. For note repertoire, higher predictor values indicate greater complexity, whereas for song linearity, lower predictor values indicate greater complexity. Estimates in the bottom row show the effect of increasing predictor values on the probability of response (vs. no response at all). Negative values indicate a greater probability of response with increasing predictor values. There was no evidence of zero inflation in the model for Fluffs, so no effects on probability of response are reported. For ease of understanding, the relationship between model coefficients and song complexity has been interpreted in the ‘Direction of effect’ column.

|  | Estimate | p-value | Direction of effect |
| --- | --- | --- | --- |
| **Number of Calls** |  |  |  |
| Note Repertoire | 0.099<br>**-0.453** | 0.114<br>**0.032** | **Birds were more likely to respond to more complex songs** |
| Song Linearity | 1.932<br>**15.477** | 0.266<br>**0.026** | **Birds were more likely to respond to more complex songs** |
| **Stimulus Order** | **-0.179** | **<0.001** | **Birds responded less to stimuli later in the trial (habituation)** |
| **Test Order** | **0.073** | **<0.001** | **Birds responded more in later trials (acceleration)** |
| **Hand-reared** | **-1.517** | **0.015** | **Hand-reared birds responded less to stimuli** |
| **Average duration** | **0.136** | **0.025** | **Birds responded more to longer songs (greater sound density)** |
| **Number of Hops** |  |  |  |
| Repertoire | 0.050<br>-0.126 | 0.533<br>0.222 | Not significant |
| *Linearity* | 3.149<br>5.643 | 0.154<br>0.076 | *Birds were more likely to respond to more complex songs* |
| **Stimulus Order** | **-0.064** | **<0.001** | **Birds responded less to stimuli later in the trial (habituation)** |
| **Test Order** | **0.164** | **<0.001** | **Birds responded more in later trials (acceleration)** |
| Hand-reared | 0.561 | 0.574 | Not significant |
| Average duration | -0.029 | 0.718 | Not significant |
| **Number of Bill Wipes** |  |  |  |
| Repertoire | 0.055<br>-0.140 | 0.539<br>0.286 | Not significant |
| Linearity | -0.128<br>7.352 | 0.962<br>0.125 | Not significant |
| **Stimulus Order** | **-0.124** | **0.035** | **Birds responded less to stimuli later in the trial (habituation)** |
| **Test Order** | **0.306** | **0.001** | **Birds responded more in later trials (acceleration)** |
| Hand-reared | -0.218 | 0.893 | Not significant |
| Average duration | -0.054 | 0.561 | Not significant |
| **Number of Fluffs** |  |  |  |
| **Repertoire** | **0.140** | **0.043** | **Birds responded more to more complex songs** |
| Linearity | -0.542 | 0.779 | Not significant |
| Stimulus Order | -0.033 | 0.620 | Not significant |
| **Test Order** | **0.167** | **0.022** | **Birds responded more in later trials (acceleration)** |
| Hand-reared | 0.011 | 0.987 | Not significant |
| Average duration | 0.050 | 0.476 | Not significant |

As in the familiarity tests, birds habituated to stimulus presentation within 10-minute trials (Table 2). There was a significant decrease in calling, hopping and bill wiping behaviour over time within trials, although there was no change in fluffing behaviour. Across trials, we found evidence for an acceleration effect for all behaviours. Birds responded more in trials later in the experiment (Table 2).

We found no clear evidence for an effect of familiarity or relatedness on responses during complexity tests. For some behaviours, birds responded differently to related compared to unrelated songs, but patterns were not consistent across tests. For most behaviours, hand-reared birds did not significantly differ from parent-reared birds in their responses. However, hand-reared individuals produced significantly fewer calls over the course of the experiment (Table 2).

## Discussion

Java sparrow females showed different behavioural reactions based on both familiarity and complexity in male songs. Females preferred their fathers’ songs over non-familial (unfamiliar) songs. Females also preferred songs with higher note repertoires, and showed a less pronounced preference for structural complexity measured as linearity. Overall, females responded more strongly to stimuli made up of longer songs, which resulted in greater sound density. Whilst yet to be confirmed in Java sparrows, the characteristics that females used to evaluate songs in our experiments may offer different information about the males that produced those songs.

Preferences for fathers’ songs are found in other, closely-related species (Fujii et al., 2021; Fujii & Okanoya, 2022; Kato et al., 2010; Riebel, 2000) and have been reported in studies using operant tasks (Kato et al., 2010; Riebel, 2000) and in those measuring behavioural responses, including copulation solicitation displays (Fujii et al., 2021; Fujii & Okanoya, 2022). Although songs can be used in a variety of social contexts, including social cohesion in estrildid finches (Loning et al., 2023), songs used exclusively in courtship displays in Java sparrows (Soma & Mori, 2015), and the behaviours observed are linked to courtship and sexual preferences in this and other estrildid finch species (Dunning et al., 2014; Soma & Iwama, 2017; Vyas et al., 2009). Whilst we did not find a significant preference for non-father familial songs, this does not necessarily mean that Java sparrow females are unable to generalize their preferences. Birds from different social lineages in our study population were not audially isolated during early development or later in life, so females may have heard song types from other social lineages, even though they did not interact with these individuals. New work is needed to determine if Java sparrows can distinguish between familiar songs (i.e., those heard during development) and other, entirely unfamiliar conspecific songs from different populations. For most behaviours measured in our study, the direction of effect was towards a stronger response to familial non-father songs compared to unfamiliar, non-familial songs, so we cannot rule out that females weakly prefer songs from their own social line. Our experiments juxtaposed fathers’ and non-familial songs in the same trials, and non-father familial and non-familiar songs in the same trials, but did not directly juxtapose fathers’ and non-father familial birds’ songs in the same trials. Thus, we have limited power to compare preferences for fathers’ and other familial birds’ songs directly.

Java sparrow females showed preferences based on song complexity. They responded more frequently and more strongly to songs with higher note repertoires. Presently, there are no reported correlations between song complexity and aspects of male quality in Java sparrows, the drivers behind these preferences remain unclear. However, preferences for large note repertoires have been reported in other estrildid finch species. In zebra finches, songs with larger syllable repertoires were preferred in tests using both phonotaxis and operant tasks (Holveck & Riebel, 2007). A similar preference for larger repertoires was found in another study examining behavioural responses to stimulus playback across females with different early acoustic experiences (Vyas et al., 2009).

Although there was no effect of song linearity on the number of calls they produced, females were more likely to call in response to more syntactically complex songs. In other estrildid species, there is some evidence for song preferences relating to syntactical complexity. Bengalese finch females stimulated with syntactically complex songs carried more nesting material than those stimulated with simple songs (Okanoya, 2012)(). In an operant task, four of eight females preferred complex songs, suggesting that syntactical complexity may play a role in shaping female preferences (Morisaka et al., 2008). However, there was clear inter-individual variation, with half of the females taking part showing no or reversed song preference. In a meta-analysis of the effects of song complexity on male reproductive success, song versatility, which is associated with structural complexity, was less correlated with reproductive success than song repertoire (Soma & Garamszegi, 2011), suggesting that structural complexity may be a less important than repertoire in shaping mate choice.

We did not systematically juxtapose short and long duration songs in our experiments, but we found evidence in both our familiarity and complexity trials that Java sparrow females prefer longer duration songs, which, due to experimental design, were also associated with higher sound density. Therefore, we cannot confirm whether females’ preference was for song length or sound density. Preferences based on song duration have been reported in a number of passerine species (Nolan & Hill, 2004)(Nolan & Hill, 2004). Song duration could provide information on current male quality to females. For example, in barn swallows (*Hirundo rustica*), song duration was negatively correlated with parasite load and males with longer songs had higher pairing success, suggesting that females may use duration to identify parasite free males (Garamszegi et al., 2005). In addition, songs require energetic costs for production, so a greater proportion of time spent singing (i.e., higher song density) may indicate higher quality males (Gil & Gahr, 2002; Nowicki & Searcy, 2004). In white-throated sparrows (*Zonotrichia albicollis*), females preferred longer songs compared to shorter songs when presented at the same rate, but when stimuli were controlled for number of notes per minute, females did not show a significant preference for longer over shorter songs (Wasserman & Cigliano, 1991). Specific preferences for song duration, rate, and output in Java sparrows could be examined through paired stimuli to disentangle the influence of song duration and song density on female preference.

Female song preferences can have implications for male reproductive success, and may result in changes in songs over time or contribute to assortative mating within or between populations. Preferences for familiar songs in a mate choice context can assist in the correct choosing of subspecies or local populations (Slabbekoorn & Smith, 2002). Strong preferences for familiar songs, and in particular songs similar to those of the father, may contribute to the formation of pre-mating reproductive barriers (Gilman & Kozak, 2015), or contribute to inbreeding through the promotion of assortative mating based on social (genetic) background (Soma, 2011). Assortative mating based on familiarity could also negatively affect population management strategies that bring birds from multiple populations together, for example, during conservation breeding programmes (Lewis, Williams, et al., 2021). Preference for song complexity during mate choice may result in improved reproductive success for males with complex songs (Soma & Garamszegi, 2011). This may drive changes in song complexity over time due to sexual selection followed by song learning in offspring. In Bengalese finches, it has been suggested that a release from predation pressure, coupled with a female preference for complex songs, has driven increases in song complexity during the domestication process (Okanoya, 2012).

Mate choice in natural situations may not always reflect preference in controlled settings. Other factors, such as additional song parameters (Dunning et al., 2020; Nolan & Hill, 2004) and male traits (Hasegawa et al., 2011) may influence pairing. However, there is evidence that preference for familiar or complex songs may affect real-world pairings. In zebra finches, females preferentially paired with males whose songs matched their populations of rearing and that more closely resembled songs of their adolescent peers when allowed to select their own mates (Wang et al., 2022). Our results show that Java sparrows, like zebra finches, show preferences for familiar songs, so it is possible that similar patterns of mate choice could occur. For song complexity, although a meta-analysis revealed a small but significant link between song complexity and male reproductive success, the effect of complexity was reduced when considering genetic reproductive success (i.e., the number of genetically related offspring produced) rather than social measures (e.g., number of females acquired, number of offspring produced within a pair) (Soma & Garamszegi, 2011). This suggests that, in natural settings, females may show different song preferences for social mates and extra-pair mating, making it difficult to predict the real-world effect of preferences measured in controlled conditions. Different preferences for social and genetic mates may also have implications for evolution: reproductive success will affect genetic change over time, but where songs are socially inherited, success in rearing social offspring is more relevant to song evolution. Java sparrow females’ preferences for song complexity may result in males with larger note repertoires having greater reproductive success, and may result in sexual selection for complex songs over time. Even if females do not show a preference when pairing, investment in breeding may be reduced with non-preferred males. In Bengalese finches, females produced heavier eggs and showed a trend towards male-biased broods when their mate had a longer song duration (Soma & Okanoya, 2013). Across tests, female Java sparrows varied in their preference and strength of responses. Therefore, individual differences may also contribute to mate choice and investment in real-world scenarios. Further investigation of mate choice in natural conditions is necessary to understand the implications of female preference for assortative mating and sexual selection, and whether this could contribute to pre-mating barriers between populations or drive vocal change across generations.

In summary, Java sparrow females showed clear preferences for specific song characteristics across the experimental tests. This suggests that overall song preference in this species is likely to be multi- faceted, with multiple song features contributing different information about the singers. In this study, we identified song familiarity, complexity, and song duration as predictors of female preference. However, other song features not tested here may also contribute to females’ overall preferences. As natural songs vary on multiple axes, and these axes may covary, it can be difficult to determine which specific features are more strongly preferred. Artificial song stimuli that differ in only one characteristic at a time may be useful in disentangling female preferences for specific aspects of song and allow us to build a more comprehensive overview of female preference as a whole.

## Author contributions

RL and MS conceptualized the study and were responsible for the experimental design. Data collection and extraction of behavioural data from videos was carried out by HM and RL. Data analysis was conducted by RL, MS, and TG. The initial draft of the manuscript was written by RL, with RL, MS, and TG contributing to reviewing and editing. MS and TG provided supervision throughout the project.

## Acknowledgements

RL’s work on this project was funded by the Natural Environment Research Council (NERC) EAO Doctoral Training Partnership (grant NE/L002469/1) in partnership with Chester Zoo, UK. RL also received funding through the JSPS Summer Programme. This project also received funding from the Daiwa Foundation for Anglo-Japanese Exchange and JSPS Grants-in-Aid for Young Scientists (grant 23680027 and 16H06177). We thank Nagisa Matsuda, Ken Otsuka, Hiroko Kagawa, and Nao Ota for archival song recordings.

## Supplementary Material 1: Results of models for familiarity with trials involving clutch mates (n=4) removed

**Table S1.1:** Results of mixed effect models for behavioural responses to song familiarity using data from stimulus presentation only. Estimates show the effect of the predictor on the logged number of responses.

|  | Estimate | p-value | Direction of effect |
| --- | --- | --- | --- |
| **Number of Calls\*** |  |  |  |
| **Father vs. Non-familial** | 0.714 | <0.001 | **Females respond more to father's calls** |
| Familial vs. Non-familial | 0.603 | <0.001 | **Females respond more to familial calls** |
| **Stimulus order** | **-0.209** | **<0.001** | **Birds responded less to stimuli later in the trial (habituation)** |
| **Test Order** | **0.111** | **<0.001** | **Birds responded more in later trials (acceleration)** |
| **Average Duration** | **0.175** | **<0.001** | **Birds responded more to longer songs (greater sound density)** |
| **Number of Hops** |  |  |  |
| **Father vs. Non-familial** | **1.930** | **<0.001** | **Females respond more to father's calls** |
| Familial vs. Non-familial | **0.784** | **0.010** | **Females respond more to familial calls** |
| **Stimulus order** | **-0.087** | **<0.001** | **Birds responded less to stimuli later in the trial (habituation)** |
| **Test Order** | **-0.110** | **<0.001** | **Birds responded less in later trials (habituation)** |
| Average Duration | 0.047 | 0.632 | Not significant |
| **Number of Bill Wipes** |  |  |  |
| Father vs. Non-familial | 0.333 | 0.634 | Not significant |
| Familial vs. Non-familial | -0.390 | 0.488 | Not significant |
| Stimulus order | 0.088 | 0.313 | Not significant |
| **Test Order** | **0.352** | **<0.001** | **Birds responded more in later trials (acceleration)** |
| Average Duration | 0.125 | 0.491 | Not significant |
| **Number of Fluffs** |  |  |  |
| **Father vs. Non-familial** | *0.763* | *0.086* | *Females respond more to father's calls* |
| Familial vs. Non-familial | 0.069 | 0.853 | Not significant |
| **Stimulus order** | **-0.275** | **0.019** | **Birds responded less to stimuli later in the trial (habituation)** |
| Test Order | 0.022 | 0.856 | Not significant |
| **Average Duration** | **0.251** | **0.022** | **Birds responded more to longer songs (greater sound density)** |

\*Model was run with a reduced zero inflation term containing an intercept only for convergence

## Supplementary Information 2: Results of models using combined behavioural data from stimulus presentation and the adjacent silent gap

The statistical models used to analyse the combined behavioural data from stimulus presentation and the adjacent silent gap follow the same format as those described in the ‘Data Analysis’ section.

**Table S2.1:** Results of mixed effect models for behavioural responses to song familiarity using combined data from stimulus presentation and adjacent silent gap. Estimates show the effect of the predictor on the logged number of responses. For bill wiping behaviour, zero-inflation was detected in the combined behavioural data, even though it was not for behavioural data in the stimulus presentation, so a zero-inflation term was included in the model.

|  | Estimate | p-value | Direction of effect |
| --- | --- | --- | --- |
| **Number of Calls\*** |  |  |  |
| **Father vs. Non-familial** | **0.723** | **<0.001** | **Females respond more to father's calls** |
| Familial vs. Non-familial | **0.588** | **<0.001** | **Females respond more to familial calls** |
| **Stimulus order** | **-0.220** | **<0.001** | **Birds responded less to stimuli later in the trial (habituation)** |
| **Test Order** | **0.112** | **<0.001** | **Birds responded more in later trials (acceleration)** |
| **Average Duration** | **0.164** | **<0.001** | **Birds responded more to longer songs (greater sound density)** |
| **Number of Hops** |  |  |  |
| **Father vs. Non-familial** | **1.409** | **<0.001** | **Females respond more to father's calls** |
| Familial vs. Non-familial | 0.427 | 0.174 | Not significant |
| **Stimulus order** | **-0.098** | **<0.001** | **Birds responded less to stimuli later in the trial (habituation)** |
| **Test Order** | **-0.115** | **<0.001** | **Birds responded less in later trials (habituation)** |
| Average Duration | -0.053 | 0.602 | Not significant |
| **Number of Bill Wipes** |  |  |  |
| Father vs. Non-familial | 0.421 | 0.532 | Not significant |
| Familial vs. Non-familial | -0.400 | 0.399 | Not significant |
| Stimulus order | -0.034 | 0.691 | Not significant |
| **Test Order** | **0.254** | **0.009** | **Birds responded more in later trials (acceleration)** |
| Average Duration | 0.077 | 0.648 | Not significant |
| **Number of Fluffs** |  |  |  |
| **Father vs. Non-familial** | **0.770** | **0.044** | **Females respond more to father's calls** |
| Familial vs. Non-familial | 0.134 | 0.696 | Not significant |
| **Stimulus order** | **-0.273** | **0.009** | **Birds responded less to stimuli later in the trial (habituation)** |
| Test Order | 0.007 | 0.946 | Not significant |
| **Average Duration** | **0.202** | **0.044** | **Birds responded more to longer songs (greater sound density)** |

\* Stepwise removal of random effects resulted in predictors explaining >1% of variance in the model being removed. The final model included a random effect of female in the conditional and zero-inflation models.

**Table S2.2:** Results of mixed effect models for behavioural responses to song complexity using combined data from stimulus presentation and adjacent silent gap. For note repertoire, higher predictor values indicate greater complexity, whereas for song linearity, lower predictor values indicate greater complexity. Estimates in the bottom row show the effect of increasing predictor values on the probability of response (vs. no response at all). Negative values indicate a greater probability of response with increasing predictor values. There was no evidence of zero inflation in the model for Fluffs, so no effects on probability of response are reported. For ease of understanding, the relationship between model coefficients and song complexity has been interpreted in the ‘Direction of effect’ column.

|  | Estimate | p-value | Direction of effect |
| --- | --- | --- | --- |
| **Number of Calls** |  |  |  |
| Note Repertoire | **0.085**<br>**-0.570** | **<0.001**<br>**0.011** | **Birds were more likely to respond and responded more to more complex songs** |
| Song Linearity | -0.591<br>10.446 | 0.179<br>0.093 | *Birds were more likely to respond to more complex songs* |
| **Stimulus Order** | **-0.164** | **<0.001** | **Birds responded less to stimuli later in the trial (habituation)** |
| **Test Order** | **0.053** | **<0.001** | **Birds responded more in later trials (acceleration)** |
| **Hand-reared** | **-1.545** | **0.016** | **Hand-reared birds responded less to stimuli** |
| **Average duration** | **0.126** | **<0.001** | **Birds responded more to longer songs (greater sound density)** |
| **Number of Hops** |  |  |  |
| Repertoire | 0.072<br>-0.084 | 0.335<br>0.378 | Not significant |
| Linearity | 1.826<br>4.525 | 0.379<br>0.149 | Not significant |
| **Stimulus Order** | **-0.064** | **<0.001** | **Birds responded less to stimuli later in the trial (habituation)** |
| **Test Order** | **0.169** | **<0.001** | **Birds responded more in later trials (acceleration)** |
| Hand-reared | 0.696 | 0.502 | Not significant |
| Average duration | -0.055 | 0.467 | Not significant |
| **Number of Bill Wipes** |  |  |  |
| Repertoire | -0.022<br>-0.109 | 0.736<br>0.373 | Not significant |
| Linearity | -0.239<br>4.403 | 0.897<br>0.236 | Not significant |
| **Stimulus Order** | **-0.162** | **0.002** | **Birds responded less to stimuli later in the trial (habituation)** |
| Test Order | 0.087 | 0.221 | Not significant |
| Hand-reared | -0.185 | 0.510 | Not significant |
| Average duration | 0.018 | 0.789 | Not significant |
| **Number of Fluffs** |  |  |  |
| **Repertoire** | *0.131* | *0.059* | *Birds responded more to more complex songs* |
| Linearity | -1.701 | 0.383 | Not significant |
| Stimulus Order | -0.014 | 0.831 | Not significant |
| **Test Order** | **0.210** | **0.002** | **Birds responded more in later trials (acceleration)** |
| Hand-reared | -0.095 | 0.881 | Not significant |
| Average duration | 0.015 | 0.828 | Not significant |

## Supplementary Material 3

**Supplementary Table 3A.** Collinearity between behavioural responses across females in familiarity trials.

|  | **Calls** | **Hops** | **Bill Wipes** | **Fluffs** |
| --- | --- | --- | --- | --- |
| **Calls** | 1.000 | 0.786 | 0.248 | 0.330 |
| **Hops** | 0.786 | 1.000 | 0.231 | 0.330 |
| **Bill Wipes** | 0.248 | 0.231 | 1.000 | 0.234 |
| **Fluffs** | 0.330 | 0.330 | 0.234 | 1.000 |

**Supplementary Table 3B.** Collinearity between behavioural responses across females in complexity trials.

|  | **Calls** | **Hops** | **Bill Wipes** | **Fluffs** |
| --- | --- | --- | --- | --- |
| **Calls** | 1.000 | 0.363 | 0.381 | 0.269 |
| **Hops** | 0.363 | 1.000 | 0.370 | 0.092 |
| **Bill Wipes** | 0.381 | 0.370 | 1.000 | 0.169 |
| **Fluffs** | 0.269 | 0.092 | 0.169 | 1.000 |

